# The Genetic Architecture of Morphological Scaling

**DOI:** 10.1101/2022.06.07.495193

**Authors:** Austin S. Wilcox, Isabelle M. Vea, W. Anthony Frankino, Alexander W. Shingleton

## Abstract

Morphological scaling relationships between the sizes of individual traits and the body captures the characteristic shape of a species, and the evolution of scaling is the primary mechanism of morphological diversification. However, we have almost no knowledge of the genetic architecture of scaling, critical if we are to understand how scaling evolves. Here we explore the genetic architecture of *population-level* morphological scaling relationships – the scaling relationship fit to multiple genetically-distinct individuals in a population – by describing the distribution of *individual* scaling relationships – genotype-specific scaling relationships that are unseen or cryptic. These individual scaling relationships harbor the genetic variation that determines relative trait growth within individuals, and theoretical studies suggest that their distribution dictates how the population scaling relationship will respond to selection. Using variation in nutrition to generate size variation within 197 isogenic lineages of *Drosophila melanogaster*, we reveal extensive variation in the slopes of the wing-body and leg-body scaling relationships among individual genotypes. This genetic variation reflects variation in the nutritionally-induced size plasticity of the wing, leg and body. Surprisingly, we find that variation in the slope of individual scaling relationships primarily results from variation in nutritionally-induced plasticity of body size, not leg or wing size. These data allow us to predict how different selection regimes alter scaling in *Drosophila* and is the first step in identifying the genetic targets of such selection. More generally, our approach provides a framework for understanding the genetic architecture of scaling, an important prerequisite to explaining how selection changes scaling and morphology.

## Introduction

Static morphological scaling relationships (commonly referred to as *static allometries* (Klingenberg and Zimmermann, 1992) describe the size relationships among morphological traits as they co-vary with body size among individuals at the same developmental stage in a population, species, or other biological group (Shingleton, 2010). In as much as the shape of an animal is determined by the relative size of its constituent body parts, differences in morphological scaling relationships capture variation in body shape within and among animal groups. Because morphological diversity is dominated by variation in animal shape, the study of morphological scaling relationships has been the focus of evolutionary biologists for well over a century (Huxley, 1924, 1932; Gould, 1966; Thompson and Bonner, 1992; Gayon, 2000). Nevertheless, until recently, almost nothing was known regarding the developmental-genetic mechanisms that regulate morphological scaling and that are the proximate targets of selection for morphological change (Tang *et al*., 2011; Emlen *et al*., 2012; Casasa *et al*., 2017). Even less is known of the distributions of genetic variation in these mechanisms that should determine how scaling responds to selection. This is primarily because, unlike most other morphological traits, scaling is ostensibly a characteristic of a group rather than an individual. Because groups of animals are, typically, genetically heterogeneous, describing the genetic architecture of morphological scaling is therefore challenging.

Historically, the literature has been concerned mostly with *population* scaling relationships (Huxley and Tessier, 1936; Gould, 1973; Klingenberg and Zimmermann, 1992; Wilkinson, 1993; Dreyer *et al*., 2016). Here, size variation among individuals results from genetic and environmental variation, and the line fit to trait-body size data reveals how these covary in a that population and environment. (For clarity, here we restrict the term ‘trait’ to morphological characteristics other than body size). When trait and body size are plotted on a log-log scale, the slope of their relationship is referred to as the *allometric coefficient* (Huxley and Tessier, 1936) which reflects the relative sensitivity of the trait and body to the myriad environmental and genetic factors that affect their size (Shingleton *et al*., 2007).

More recently, attention has turned to *individual* scaling relationships (Figure 1; Dreyer et al. 2016). These result from co-variation in trait and body size due to variation in a single environmental or genetic factor, with all other size regulatory factors held constant (including genotype). Variation in slopes and intercepts among individual scaling relationships reflect genetically-based differences among individuals in how trait and body size respond to the varying size-regulatory factor. When the size-regulatory factor is environmental – yielding an individual environmental scaling relationship – variation among scaling relationships is a consequence of genotype-by-environment interactions. Importantly, different environmental factors will generate different scaling relationships for the same genotype. For example, in *Drosophila melanogaster*, the effects of nutrition during development on body size generate different morphological scaling relationships among traits than the effects of temperature during development (Shingleton *et al*., 2009). Similarly, when the size-regulatory factor is genetic – yielding an individual genetic scaling relationship – variation among scaling relationships is a consequence of genotype-by-genotype interactions. This would be generated by allelic variation at a single locus interacting epistatically with an otherwise constant genetic background. In reality each individual occupies only a single point on their individual scaling relationships, reflecting the particular combination of environmental and genetic effects that determine trait and body size in that individual. Fitting a line to a collection of these points from genetically heterogenous individuals in a population, each experiencing a unique combination of environmental factors, generates a population scaling relationship (Figure 1A).

**Figure 1:**
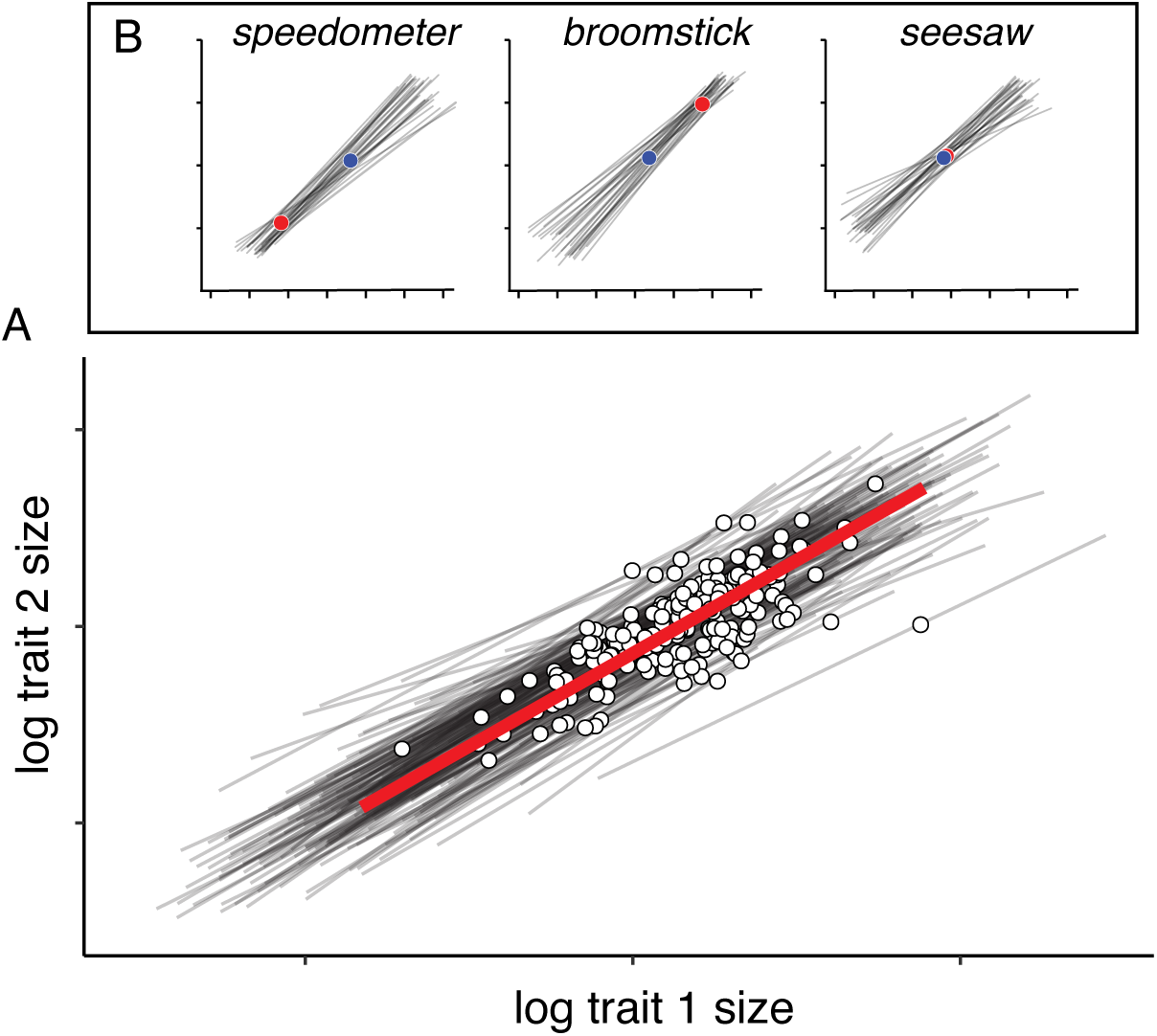
Individual and population scaling relationships. (**A**) Individual scaling relationships (thin grey lines) result from variation in body size due to variation in a single environmental or genetic factor, with all other size regulatory factors held constant. However, because each individual has a single genotype and is exposed to a single combination of environmental factors, it occupies only a single point on its individual scaling relationship (white circles). The observed population scaling relationship (red line) is the scaling relationship among individuals in a population. (**B**) The distribution of individual scaling relationships determines how the population scaling relationship responds to selection (Frankino *et al*., 2019), and can be *speedometer*, *broomstick*, or *seesaw*, depending on where the median point of intersection (red circle) lies relative to the bivariate mean of trait sizes (blue circle).

Thus, underlying population scaling relationships are collections of unseen, or cryptic, individual scaling relationships. The distribution of individual scaling relationships in a population will place individuals at particular locations around the population scaling relationship, and selection on these individuals may alter the distribution of individual scaling relationships, which in turn may change the slope or intercept of the population-level scaling relationship (Dreyer *et al*., 2016; O’Brien *et al*., 2017; Houle *et al*., 2019). Mathematical modeling suggests that the response of the population scaling relationship to selection is dependent on the distribution of the underlying individual scaling relationships; the same selective pressure applied to two ostensibly identical population scaling relationships can generate very different responses if the underlying distribution of individual scaling relationships differs (Dreyer *et al*., 2016). Consequently, if we are to understand the evolution of population scaling relationships, we need to understand the genetic architecture of the individual scaling relationships that underlie them.

While the concept of individual scaling relationships is straightforward, measuring them is not. Individual environmental scaling relationships can be generated by fitting a line to the trait-body size combinations expressed by genetically-identical individuals reared across an environmental gradient. Individual genetic scaling relationships can be generated by fitting a line to the trait-body size combinations expressed by individuals possessing allelic variation at only a single locus in an otherwise co-isogenic background and reared in a single environment. For many animals, such environmental and genetic control is impractical or impossible to impose. The measurement of individual scaling relationships is tractable in *Drosophila*, however, as the long-term maintenance of (near) isogenic populations is routine and genetic variation can be generated at a single gene or locus (Frankino *et al*., 2019; Houle *et al*., 2019).

In this paper we focus on the genetic architecture of population scaling relationships by characterizing the distribution of individual scaling relationships, using isogenic lineages of *D. melanogaster* as a model. The individual scaling relationships for each genotype were generated by varying access to food during development; because trait and body size results from variation in developmental nutrition, we refer to these individual scaling relationships as *nutritional scaling relationships* (Dreyer *et al*., 2016). We have previously used this simple diet manipulation to generate variation in wing and body size in *D. melanogaster* (Stillwell *et al*., 2011; Frankino *et al*., 2019). Here we apply this approach to 197 isogenic lineages of *D. melanogaster*, to determine the distribution of individual nutritional wing- and leg-body scaling relationships in this population. Further, we assay the nutritionally-induced size plasticity of these traits and the body within each lineage. We use these data to explore the genetic architecture of nutritional scaling within and among traits and the variation in relative trait plasticity that accounts for this architecture.

## Material and Methods

### Fly Stocks

All flies used in this study came from The *Drosophila* Genome Resource Panel (DGRP). The DGRP is a library of ∼200 fully sequenced inbred isogenic *Drosophila* lineages that originated from a single outbred population (Mackay *et al*., 2012) collected from Raleigh, NC, USA. Flies were maintained on standard cornmeal molasses medium (Frankino *et al*., 2019) and maintained on a 12:12 light cycle at 22°C and 75% humidity.

### Starvation treatment

*Drosophila* egg collection, rearing, and phenotyping followed our established protocols (Stillwell *et al*., 2011, 2016; Frankino *et al*., 2019). For each DGRP lineage, females oviposited for three days. At 24h, 48h and 72h, eggs were collected, divided into lots of 50 and placed into vials containing 10ml of standard cornmeal molasses medium. This generated three age cohorts of flies (D0, D1 and D2, respectively). When third instar larvae from D0 began to pupariate, larvae from each cohort were removed from the food and placed into empty food vials with a wet cotton plug to provide moisture. Pupae were removed from these vials and transferred to individual 1.5ml Eppendorf tubes, each with a small hole in the lid, to complete development to adulthood. Larvae in the D0 cohort were starved for between 0-24h before pupariation, larvae in the D1 cohort were starved for between 24-48h before pupariation, and larvae in the D3 cohort were starved for between 48-72h before pupariation. Because larvae stop feeding ∼24h before pupariation (Testa *et al*., 2013), D0 larvae were essentially allowed to feed *ad libitum* and more-or-less achieved full adult body size. In contrast, D1 and D2 larvae were starved before larval wandering, reducing adult size depending on their size at initiation of starvation. Across all three cohorts, our starvation treatment therefore generated nutritionally-induced variation in body size. Flies were collected in nine temporal blocks, with five lineages repeated across all blocks to serve as a control.

### Body and Trait Size Measurement

Body and trait size were measured using established protocols (Shingleton *et al*., 2009; Stillwell *et al*., 2011). Briefly, *Drosophila* adults were dissected, and their right wing and right first leg mounted in dimethyl hydantoin formaldehyde (DMHF). Pupal area (a proxy for body size), wing area, and femur length (a proxy for leg length) were measured across the full range of body size for ∼50 individuals per sex per lineage; Figure 2). All traits were measured via semi-automated custom software (Metamorph, Molecular Devices LLC) that captures images from a digital camera-equipped microscope (Leica DM6000B, Leica Microsystems Inc). Femur length was squared to put it in the same dimension as wing and pupal area, and all measurements were log transformed to ensure scale invariance across traits of different sizes.

**Figure 2:**
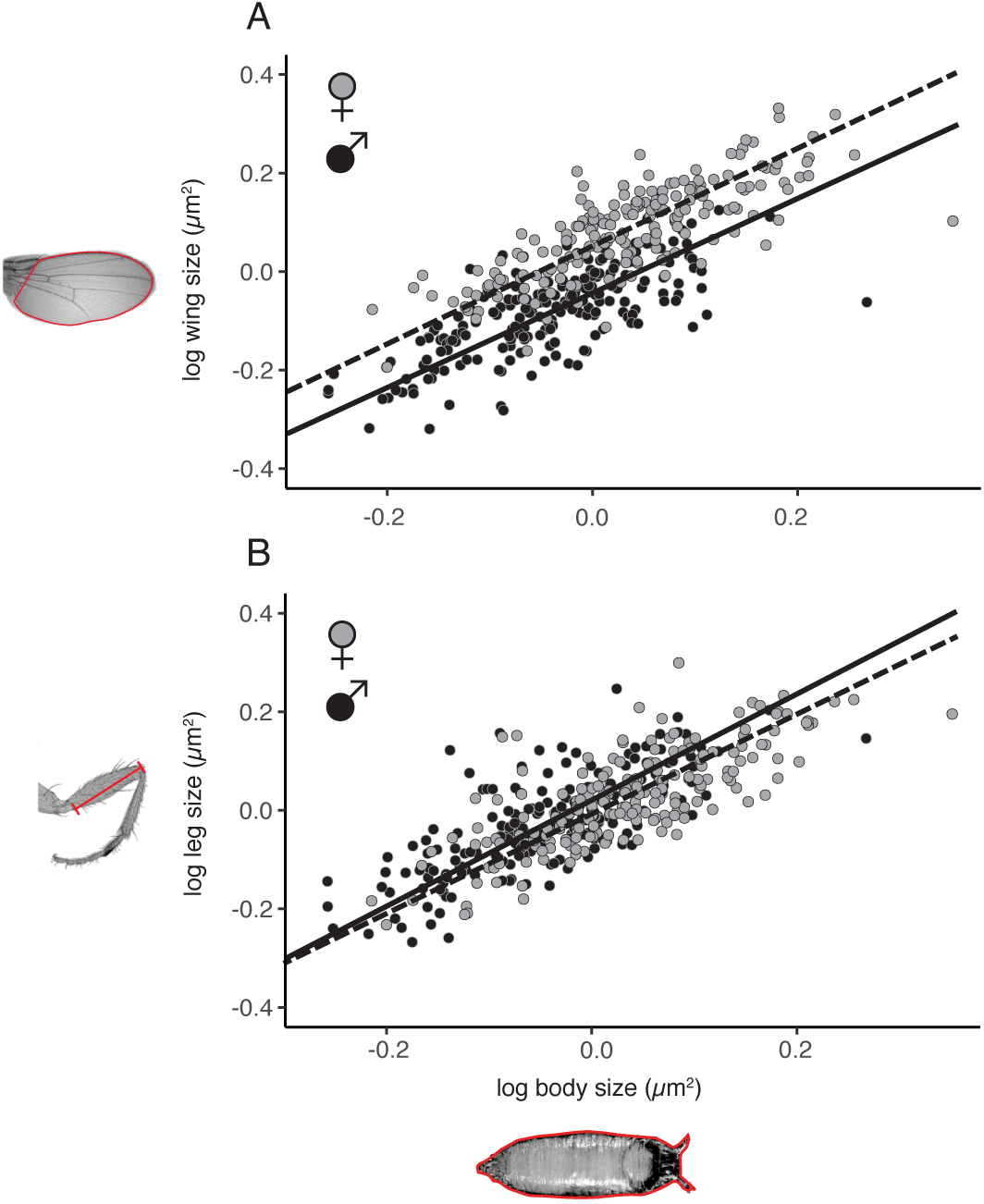
The population scaling relationships of (A) wing-body and (B) leg-body size in female (gray points, broken line) and male (black points, solid line) *Drosophila*. Points show the mean wing/leg/body size of all flies in each lineage. Lines show the mean population scaling relationship, generated by sampling a single individual from each lineage, fitting the MA regression and repeating 10,000 times by sex. For both wing-body and leg-body scaling relationships, there is a significant difference in intercept but not slope between females (gray) and males (black) (Table 1). The measurements taken are shown in red on the images of the wing, leg and pupa.

### Statistical Analysis

All data as well as the *R* scripts used to analyze them are provided on Dryad. We collected data from >12 flies per sex per lineage, with a mean of 65 flies per sex per lineage. Block effects were removed by fitting the model *T = K* to the data, where *T* is body/trait size and *K* is block. We then used the residuals of the fit for each trait/body as a measure of trait/body size independent of block. Theoretical studies indicate that major axis (MA) model II regression best captures the developmental mechanisms that generate morphological scaling relationships (Shingleton, 2019), so where possible we used this method to fit the individual scaling relationships. However, for completeness, and when testing more sophisticated models (e.g. when lineage was treated as a random factor) we fit the relationship using Model I linear regression, using maximum likelihood (R package: lme4; Bates et al. 2014), and Bayesian methods (R package: MCMCglmm; Hadfield 2010).

## Results

### Population and Individual Morphological Scaling Relationships

Almost all published scaling relationships are population-level scaling relationships, where each point on a plot of body size against trait size is a genetically distinct individual. To estimate the population scaling relationship between wing or leg and body (pupal) size in our *Drosophila* population, we first sampled one individual of each sex from each lineage (genotype), pooled these observations to create a population, and then calculated the slope and intercept of the major axis (MA) Model II regression of trait size against body size. We repeated this 10,000 times to generate a 95% confidence intervals for the slope and the intercept for the female and male wing-body and leg-body population morphological scaling relationships (Table 1, Figure 2). There were no differences between the sexes in the slope of either of these scaling relationships (Table 1). In contrast, the intercept for the wing-body size population-level scaling relationship was higher in females than in males, while the intercept for the leg-body population scaling relationship was higher in males than in females (Table 1). This was supported by an MA regression of mean trait size against mean body size among lineages (Figure 2), which also detected no sex differences in the slope of either the wing-body or leg-body population scaling relationship (wing-body slope: *P* = 0.806; leg-body slope: *P =* 0.315, *n* = 194), but found a significant sex difference in intercept (wing-body intercept & leg-body intercept: *P* < 0.0001 for both, *n* = 194).

**Table 1:**
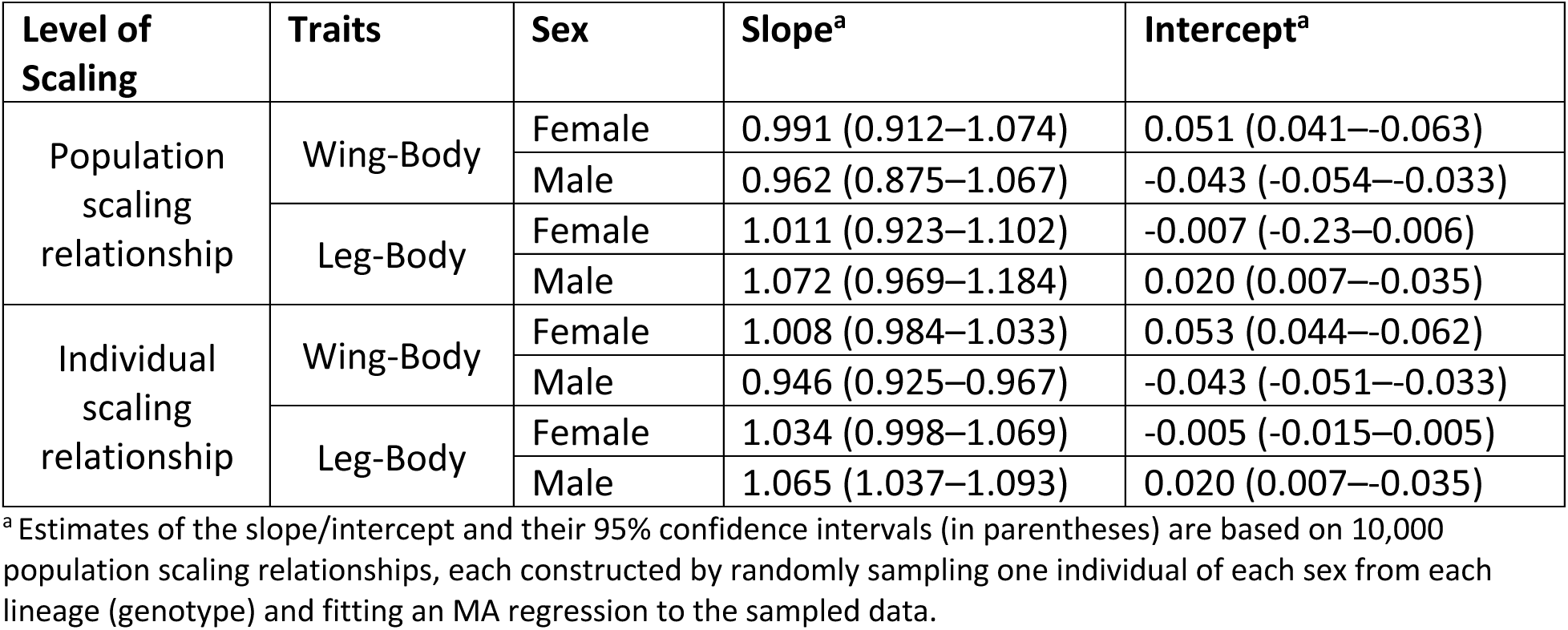
The slope and intercept of the wing-body and leg-body population and individual scaling relationships in males and females

We next explored the individual scaling relationships across the 197 isogenic lineages. We estimated the slope and intercept of the wing-body and leg-body individual scaling relationships for each sex in each lineage using MA regression. Across lineages, the wing-body individual scaling relationships were steeper and had a higher intercept in females than in males (paired t-test, *p*<0.0001 for slope and intercept, Table 1), indicating that females had larger wings than males relative to body size, and that this difference in relative wing size increased disproportionally to overall body size within lineages. In contrast, the leg-body scaling relationship had lower intercept in males than females (paired t-test, *p*<0.0001, Table 1), and tended to be shallower, although the difference in slope was not significant (paired t-test, *p*=0.0831, Table 1). These distributions were supported when fitting the individual scaling relationships using a linear-mixed model and treating lineage as a random factor (Supplementary Tables 2 and 3).

**Table 2:** Variances (diagonal), Covariances (below diagonal) and Pearson’s Correlations (above diagonal) of trait and body size plasticity within and between sexes ^a^

|  |  | male |  |  | female |  |  |
| --- | --- | --- | --- | --- | --- | --- | --- |
|  |  | wing | leg | pupa | wing | leg | pupa |
| male | wing | **0.0084** | 0.80 | 0.64 | 0.57 | 0.43 | 0.50 |
|  | leg | 0.0081 | **0.0122** | 0.64 | 0.50 | 0.48 | 0.46 |
|  | pupa | 0.0049 | 0.0059 | **0.0070** | 0.47 | 0.39 | 0.63 |
| female | wing | 0.0051 | 0.0054 | 0.0038 | **0.0095** | 0.76 | 0.63 |
|  | leg | 0.0044 | 0.0060 | 0.0037 | 0.0083 | **0.0127** | 0.64 |
|  | pupa | 0.0045 | 0.0050 | 0.0052 | 0.0061 | 0.0071 | **0.0097** |

<sup>a</sup> The darker color the higher the covariance/correlation. All correlations are significant at $P < 0.0001$ .

Within females and males, there was significant variation among genotypes in slope for both the wing-body and leg-body individual scaling relationships (Figure 2), when the relationships were fit using either an MA regression (treating lineage as a fixed factor; Supplementary Table 4) or a linear mixed-model regression (treating lineage as a random factor; Supplementary Table 5). For females, the coefficient of variation (CV) for the wing-body and leg-body MA slopes was 17.2% and 24.2% respectively, while for males the CV for the wing-body and leg-body MA slopes was 15.4% and 18.6% respectively. An important caveat is that these estimates of genetic variation are among isogenic lineages and so may not reflect the additive genetic variation for slope in an outbred population (Houle *et al*., 2019). Among lineages, there was a significant correlation between male and female slopes for both the wing-body scaling relationship (ρ = 0.30, 95% CI: 0.16-0.42) and the leg-body scaling relationship (ρ = 0.40, 95% CI: 0.28-0.52). Fitting an MA regression to this correlation revealed that, for both wing-body and leg-body scaling relationships, as the slope of the individual scaling relationship increased among lineages, the female slope increased more than the male slope (Supplementary Figure 1).

### Distribution of Cryptic Individual Scaling Relationships

Theoretical studies suggest that the distribution of individual scaling relationships in a population determines the response to selection on the scaling relationship slope (Dreyer *et al*., 2016). These distributions can be classified as either broomstick, seesaw, or speedometer; these names are derived from objects that move in a manner that looks like a plot of individual scaling relationships under each distribution. Classification is determined by where the morphological scaling relationships, on average, intersect relative to the observed range of trait sizes (Figure 3). Previously, we used the mean point-of-intersection among all pairs of individual scaling relationships to classify their distribution (Frankino *et al*., 2019). However, pairs of near-parallel individual scaling relationships can generate substantial outliers in the distribution of points-of-intersection, biasing the mean. To circumvent this problem, here we instead used the median point of intersection (MPI) to classify the distribution of the individual scaling relationships. More specifically, we compared where the MPI lies relative to the observed morphological scaling relationships (Figure 3). For a speedometer distribution, the MPI is closer to the origin than is the bivariate mean; for a broomstick distribution, the MPI is farther from the origin than is the bivariate mean; finally, for a seesaw distribution, the MPI lies approximately at the bivariate mean. For both the individual wing-body and the leg-body size scaling relationships, the MPI was close to the bivariate mean trait size in males and in females: that is, the distribution of individual scaling relationships appeared to be a seesaw in both sexes.

**Figure 3:**
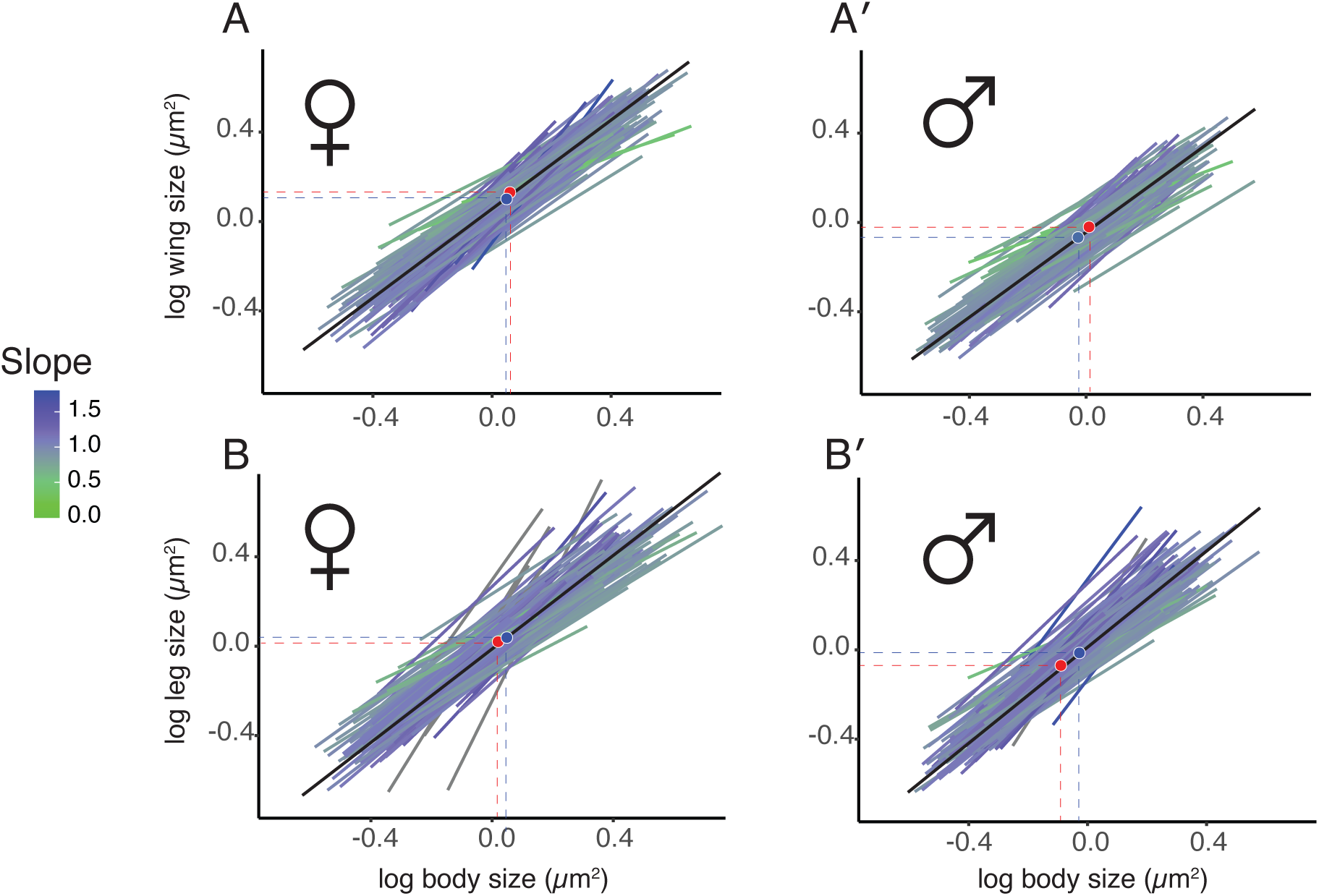
Distribution of individual scaling relationships among isogenic fly lineages. (**A, Aʹ**) The distribution of wing-body individual scaling relationships in females (A) and males (Aʹ). Males have proportionally smaller wings and shallower slopes than females (Table 1). (**B, Bʹ)** The distribution of leg-body individual scaling relationships in females (B) and males (Bʹ). Males have proportionally larger legs and steeper slopes than females (Table 1). The steepness of the slope is indicated by color (green = shallow, blue = steep). The blue circle shows the bivariate mean of trait-body size. The red circle shows the median point of intersection (MPI) for the lines. The black line is the mean individual scaling relationship across all lineages. All the scaling relationships extend two standard deviations above and below the mean body size for each lineage. Dashed lines indicate bivariate mean (blue) and MPI (red) for body size and trait size.

An artificial selection experiment to increase or decrease relative wing size (wing:thorax ratio) resulted in a corresponding increase or decrease in the slope of the wing-thorax scaling relationship, respectively (Robertson, 1962). This finding suggests a positive correlation between relative wing size and the slope of the wing-body scaling relationship among genotypes. This would occur if the distribution of individual scaling relationships were speedometer (Figure 1), which appears to contradict our observation that the distribution of wing-pupal individual scaling relationships is seesaw. Indeed, we found no correlation between a lineage’s relative wing size and the slope of its wing-body scaling relationship, in either males or females (OLS regression: *R^2^* < 0.006, *P* >0.31 for both).

However, unlike our study, Robertson (1962) did not use diet manipulation to increase the range of body size among flies, and so likely selected only well-fed individuals. In our study, these are the largest flies that occupy the upper-right portion of their individual scaling relationships (black Lines, Figure 4A). For a seesaw distribution (Figure 1B), large size-class flies will also show a positive correlation between their wing-body slope and relative wing size among lineages. This hypothesis was supported by our data. We found there was a significant positive relationship among lineages between mean relative wing size for the largest 25% of individuals in a lineage and the slope of the lineage’s wing-pupal scaling relationship, in both males and females (Figure 4B’ & C’). Conversely, for a seesaw distribution, the smallest flies should show a negative correlation between wing-body slope and relative wing size (grey lines, Figure 4A), which was also supported by our data (Figure 4B & C).

**Figure 4:**
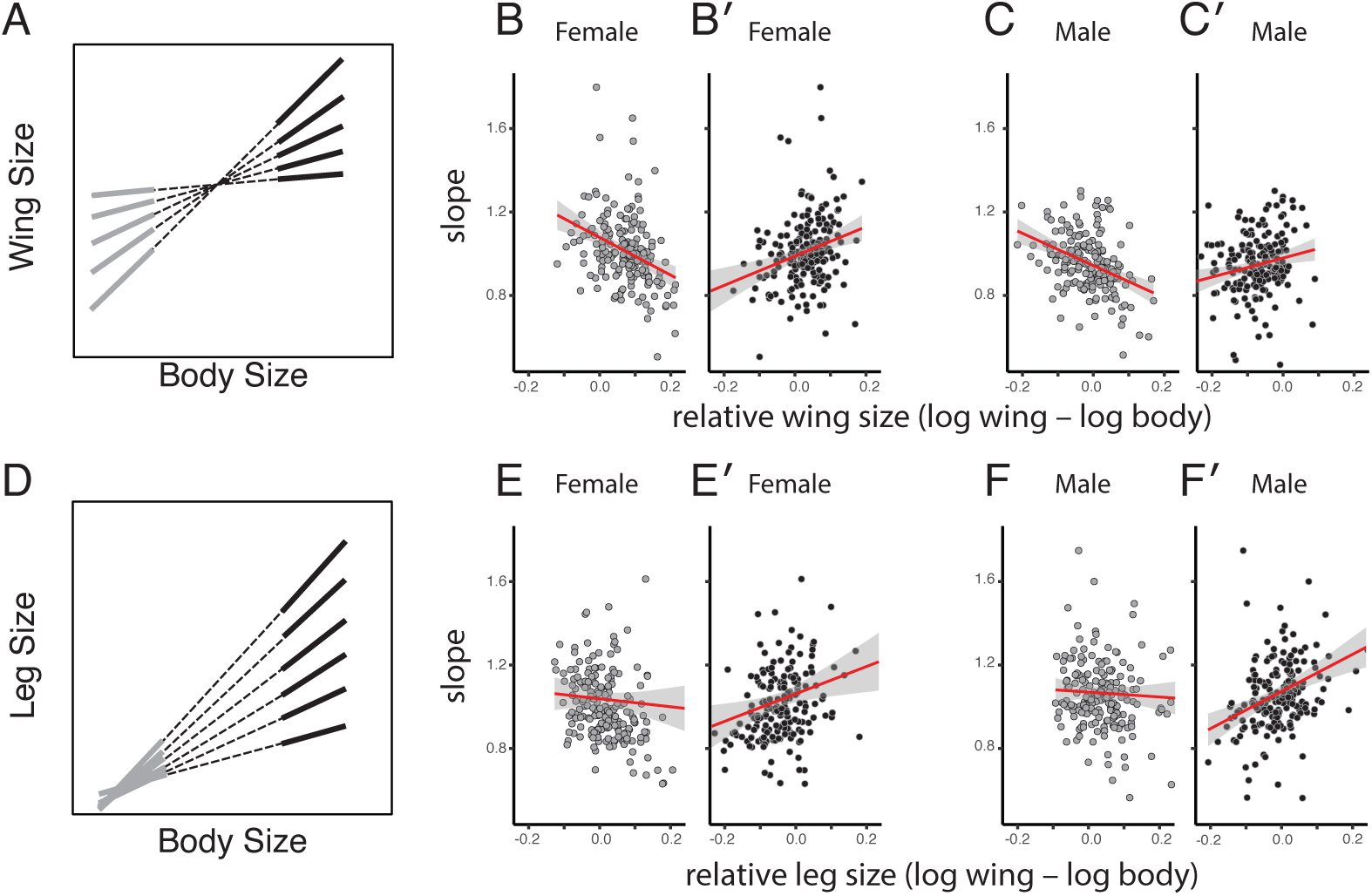
The relationship between mean relative wing/leg size and the slope of individual scaling relationships among isogenic fly lineages. (**A**) For a seesaw distribution of individual cryptic scaling relationships (dashed lines) for the largest individuals (black lines) there should be a positive correlation between their mean relative wing size in each lineage and the slope of the lineage’s individual scaling relationship, whereas this relationship should be negative among the smallest individuals (gray lines). (**B,C**) For both females (B, Bʹ) and males (C, Cʹ), there was a significant positive relationship between relative wing size for the largest 25% of individuals in each lineage, and slope of the lineage’s individual wing-body scaling relationship (OLS regression: slope = relative wing size, *F_1,192_* >8.98*, P* <0.003 for both), and a significant negative relationship between relative wing size for the smallest 25% of individuals in each lineage, and slope (OLS regression: slope = relative wing size, *F_1,192_* >26.79*, P* <0.001 for both). (**D**) For a speedometer distribution of individual scaling relationships (dashed lines), for the largest individuals (black lines) there should be a positive correlation between their mean relative wing size in each lineage and the slope of the lineage’s individual scaling relationship, whereas this relationship should be weaker or absent among the smallest individuals (gray lines). (**D, E**) For both females (D, Dʹ) and males (E, Eʹ), there was a significant positive relationship between relative leg size for the largest 25% of individuals in each lineage, and slope of the lineage’s individual leg-body scaling relationship (OLS regression: slope = relative leg size, *F_1,192_* >7.17*, P* <0.008 for both), but no relationship between relative wing size for the smallest 25% of individuals in each lineage, and slope (OLS regression: slope = relative leg size, *F_1,192_* <0.65*, P* >0.42 for both).

We also examined the relationship among lineages between the leg-body slope for a lineage and relative leg size in the largest and smallest individuals from that lineage. As was the case for the wing, there was a positive correlation between mean relative leg size for the largest 25% of individuals in a lineage and the slope of the lineage’s leg-pupal scaling relationship, in both males and females (Figure 4E’ & F’). We could not detect, however, any correlation between relative leg size and slope using data from the smallest 25% of individuals in each lineage (Figure 4E & F). This suggests that the distribution of individual scaling relationships between the leg and the body is more of a speedometer than seesaw (Figure 1B).

### Morphological Scaling and Size Plasticity

Individual scaling relationships reflect variation in body size and covariation in trait size; that is, size plasticity caused by a particular environmental or genetic factor. When size variation is due to an environmental factor, the slope of an Individual scaling relationship captures the genotype-specific size plasticity of the trait (on the *y* axis) relative to that of the body (on the *x* axis) (Shingleton *et al*., 2007). When trait size is more plastic relative to body size, the slope of the scaling relationship is greater than one; when the trait exhibits less size plasticity than the body, the slope will be less than one. Variation in the slope of individual nutritional scaling relationships can therefore be due to variation in the plasticity of trait size, variation in the plasticity of body size, or some combination of both.

To explore the relationship between trait- and body-size plasticities and the slope of individual scaling relationships, we used the range in trait and body size between the largest and smallest 10% of individuals within a lineage as a measure of size plasticity. We found significant correlations between the plasticity of trait pairs (wing v. leg, leg v. body, wing v. body) both within and between sexes (Table 4). Similarly, we also found significant correlations in the plasticity of the same trait between sexes (Table 4). We may *a priori* expect trait plasticities to be correlated due to the systemic effects of nutrition on overall body size (Shingleton *et al*., 2007). We therefore regressed wing- and leg-size plasticity against body-size plasticity using OLS regression, and used the residual values as a measure of trait-size plasticity that was independent of body-size plasticity. This analysis revealed significant correlations in trait-size plasticity among appendages and between sexes, independent of body-size plasticity (Supplementary Table 6).

Finally, we investigated the extent to which trait- or body-size plasticity explains among-genotype variation in the slope of the wing-body and leg-body individual scaling relationships. To do this, we regressed the slope of the individual scaling relationships against their trait- and body-size plasticities, by sex across all lineages. The *R^2^* of these linear regressions capture the proportion of slope variation that is due to plasticity in trait size relative to plasticity in body size (Supplementary Table 7). For both sexes, variation in the plasticity of body size explained more of the variation in the slope of the individual scaling relationships than did variation in the plasticity of either the wing or the leg (Figure 5). Indeed, there was no significant relationship between leg plasticity and the slope of the leg-body individual scaling relationship among lineages for either sex (Figure 5 C & D). Further, variation in body size plasticity explained more of the variation in the slope of individual scaling relationships in females than in males, for both wing-body and leg-body scaling. This suggests that variation in body size plasticity is greater in females than in males. Pairwise comparisons of the wing, leg and body size plasticity variances between females and males supported this hypothesis: Variance in body size plasticity was significantly greater among females than among males (*F*_192,192_ = 1.40, *P*=0.02), which was not true for variance in wing size plasticity (*F*_192,192_ = 1.13, *P*=0.37), or leg size plasticity (*F*_192,192_ = 1.05, *P*=0.74).

**Figure 5:**
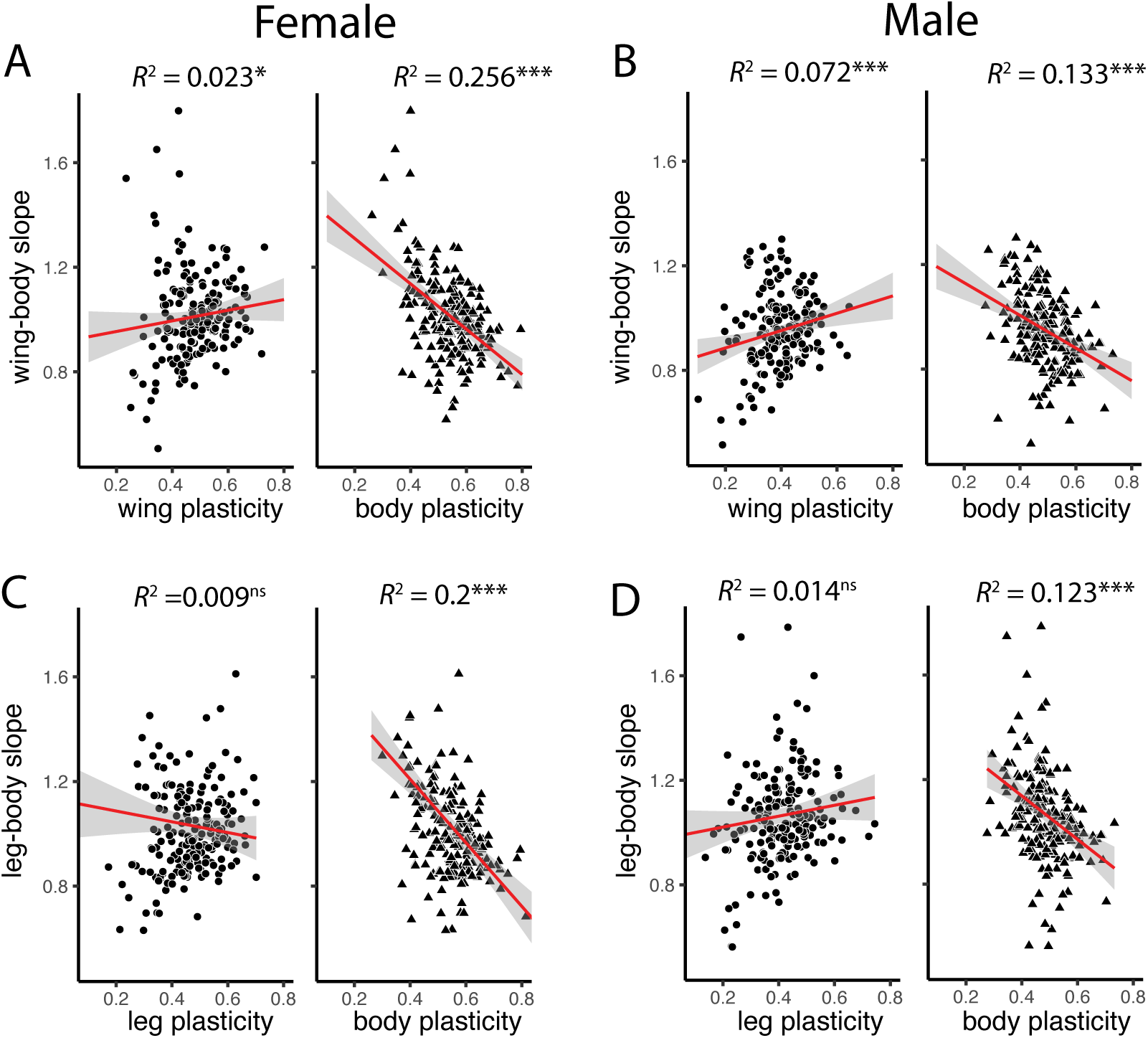
The relationship between wing-, leg-, and body-size plasticity in a lineage and the slope of the lineage’s wing-body and leg-body scaling relationship. The *R^2^* for the relationship between wing-, leg-, and body-size plasticity and the slope of the wing/leg-body size scaling relationship, among lineages, captures the proportion of variation in slope that is due to variation in wing/leg- or body-size plasticity. (**A, B**) Variation in the slope of the wing-body scaling relationship is due to variation in both wing- and body-size plasticity in both females and males, although variation in body-size plasticity is more important in males. (**C, D**) In both males and females, variation among lineages in the slopes of the leg-body individual scaling relationships is due primarily to variation in body-size plasticity. ^ns^ = non-significant, * *P*< 0.05, *** *P*<0.0001. Grey shading is 95% confidence interval of the slope. All relationships were fit using OLS linear regression.

## Discussion

Evolution of morphological scaling dominates the generation of morphological diversity among species, and yet we know little of how selection targets the developmental-genetic mechanisms that regulate trait and body size to create this diversity. Theoretical studies that distinguish between population and individual scaling relationships have hypothesized that the distribution of cryptic individual scaling relationships (seesaw versus speedometer versus broomstick) determines how the population-level scaling relationship will respond to selection (Dreyer *et al*., 2016). Our study explores variation among individual scaling relationships derived from isogenic *D. melanogaster* lineages, and how this variation relates to nutritionally induced size plasticity of two traits and the body. We find that, across the full range of body sizes generated by variation in developmental nutrition, the distribution of individual scaling relationships is approximately a seesaw for both wing-body and leg-body scaling. Further and somewhat surprisingly, we find that variation in the slope of the individual scaling relationships, which reflects the relative nutritional plasticities of trait and body size, is primarily a consequence of variation in the relative plasticity of body size. These data provide important insight into the genetic basis of variation in morphological scaling, and how this variation may respond to selection to generate morphological diversity through evolution of the population-level scaling relationship.

### Evolvability and the Genetic Architecture of Scaling

While the slopes of morphological scaling relationships can vary dramatically among species, particularly for exaggerated traits used to attract or compete for mates (Baker and Wilkinson, 2001), this seems to be the exception rather than the rule: For most species and most traits, the slopes of morphological scaling relationships tend to be evolutionarily invariant (Gould, 1966; Pelabon *et al*., 2014). These observations have led to the hypothesis that morphological scaling relationships are evolutionarily constrained by developmental or physiological mechanisms (Pelabon *et al*., 2014). Developmental studies in *Drosophila*, however, reveal that simple changes in the expression of a single gene are sufficient to substantially alter the slope of trait-body scaling for traits that otherwise maintains a near constant allometric coefficient across species, such as the wing or genitalia (Tang *et al*., 2011; Shingleton and Tang, 2012). Subsequent studies have changed gene expression to alter the slope of genital-body scaling in dung beetles (Casasa and Moczek, 2018), mandible-body scaling in male flour beetles (Okada *et al*., 2019), and horn-body scaling in rhinoceros beetles (Ohde *et al*., 2018). Consequently, it does not appear that the evolutionary invariance of morphological scaling relationship slope is rooted in developmental or physiological constraint, at least mechanistically.

An alternative explanation for the evolutionary conservation of scaling is that there is little genetic variation in the developmental mechanisms that regulate morphological scaling relationships, and upon which selection can act. This would be evident as a lack of genetic variation in the slopes of individual morphological scaling relationships within a population. Hitherto, there have been almost no data on the genetic variation of either the slope or intercept of morphological scaling (Frankino *et al*., 2019). However, our study reveals considerable variation in the slope of both the wing-body and leg-body individual scaling relationships, with coefficients of variation comparable to that for overall body size (Lafuente *et al*., 2018). A similar study on the scaling relationship between wing-vein length and wing size also revealed considerable genetic variation in the slope of individual scaling relationships (Houle *et al*., 2019). Thus, any observed inertia in the evolution of trait-body scaling relationship slope does not appear to result from lack of genetic variation, at least in *Drosophila*. An important caveat, however, is that the variation in slope detected in both this and previous studies (Frankino et al. 2019; Houle et al. 2019) is among isogenic lineages and may not reflect the level of additive genetic variation for the slope of individual scaling relationships in natural populations.

If the slopes of morphological scaling relationships are not developmentally or physiologically constrained, and if they possess levels of genetic variation comparable to that for body size, which responds rapidly to artificial selection (Hillesheim and Stearns, 1991; Partridge and Fowler, 1993; Turner *et al*., 2011), we should expect morphological scaling to also respond rapidly to artificial selection. However, artificial selection on the slope of the wing-body scaling relationship in *Drosophila* revealed an erratic and weak response, with an apparent heritability of less than 0.015 (Stillwell *et al*., 2016). The imposed selection regime attempted to alter the slope of the population wing-body scaling relationship without changing either mean wing or mean body size (i.e., to rotate the scaling relationship approximately about the bivariate mean). To increase the slope, these investigators selected large-bodied individuals with disproportionally large wings and crossing them with small-bodied individuals with disproportionally small wings. To decrease the slope, they selected large-bodied individuals with disproportionally small wings and crossing them with small-bodied individuals with disproportionally large wings. The authors ascribed the low response to pleiotropy between the slope and mean trait and body size (Stillwell *et al*., 2016). However, an alternative explanation is that the selection regime failed to consider the relationship between the observed population-level scaling relationship and the underlying distribution of individual cryptic scaling relationships. That is, the individuals selected because of their disproportionally sized wings may have possessed individual wing-body scaling relationships that would not facilitate the desired response to selection. This would occur if the slope distribution of the individual scaling relationship were of the speedometer or broomstick distribution (Dreyer *et al*., 2016). The same reasoning may explain why another selection experiment, which attempted to change the slope of the scaling relationship between wing-vein length and wing size, had a similarly weak and erratic response when the full range of body size was produced via diet manipulation (Bolstad *et al*., 2015). This latter study also ascribed the relative lack of response to pleiotropic constraints (Houle *et al*., 2019), rather than a failure of the selection regime to efficiently target alleles that regulate the slope of the scaling relationship.

Our data detailing the distribution of individual scaling relationships in a population – albeit among homozygous genotypes – will facilitate the design of artificial selection regimes that most efficiently target the slope of individual scaling relationships. The efficacy of such selection regimes will provide a nuanced method to test of the pace and extent to which the slopes of morphological scaling relationships can evolve. Indeed, earlier artificial selection experiments to shift the intercept of morphological scaling relationships in *Drosophila* (Robertson, 1962) and stalk-eyed flies (Wilkinson, 1993) – by selecting to increase relative wing size and eye-span – rapidly and indirectly altered the slope of the relationship. A third study, that applied directional selection on body size in the tobacco hornworm *Manduca sexta* also indirectly altered the slope of the wing-body scaling relationship (Tobler and Nijhout, 2010). Our data (Figure 3) may reveal why such selection will be effective, at least with respect to the wing-body scaling relationship in *Drosophila*: Selection to increase relative wing size in large well-fed flies will indirectly select to increase the slope of the wing-body scaling relationship (Figure 4 B’ and C’).

Why then, given the apparent extent of genetic variation underlying the slope of population-level morphological scaling relationships, do their slopes appear to be evolutionarily constrained? Our data, along with those of Houle et al. (Houle *et al*., 2019), support the hypothesis that the evolutionary conservatism of morphological scaling relationship slopes is a consequence of natural selection, which will favor proportions that enable ecological performance. This may be particularly true for appendages that are involved in mobility, such as wings and legs, where changes in loading – that is total body mass divided by appendage dimensions (Gilchrist and Huey, 2004; David *et al*., 2011) – may have substantial energetic or functional consequences. An alternative, non-exclusive, hypothesis is that changes in the slope of scaling relationships reduce fitness due to pleiotropic effects, for example by altering the scaling relationship between other traits and body size (Houle *et al*., 2019). While this may be the case for the scaling relationships among traits in a highly integrated organ, for example the veins of the wing (Houle *et al*., 2019), this does not appear to be true for the relationship among appendages: Developmentally altering the scaling relationship between wing and body size, for example, does not affect the scaling relationships between body size and other traits (Tang *et al*., 2011). Nevertheless, our data indicate a tight genetic correlation in size plasticity among traits independent of body size plasticity. Because linkage disequilibrium breaks down over short distances in the population of flies used in our study (Mackay *et al*., 2012), this correlation likely arises from pleiotropy, which would need to be broken for natural selection to change the slope of one trait’s morphological scaling relationship with body size independently of another. Exploring the fitness of flies that have been allometrically engineered to have atypical scaling relationships, generated using either transgenics or artificial selection, will help resolve these questions (Wilkinson and Reillo, 1994; Frankino *et al*., 2005, 2007; Houle *et al*., 2019).

### Size Plasticity and the Genetic Architecture of Scaling

The slope of nutritionally-generated individual scaling relationships reflects the relative nutritional plasticity of trait and body size (Shingleton *et al*., 2007). Variation among the slopes of these scaling relationships can result from genetic variation in relative body-size plasticity, trait-size plasticity, or both. From a developmental perspective, both trait and body size plasticity is a response to developmental nutrition, mediated through systemic growth-regulatory mechanisms, canonically the IIS and TOR signaling pathways (Vea and Shingleton, 2020). Autonomous changes in a trait’s growth-sensitivity to variation in either IIS or TOR signaling is sufficient to alter the slope of the trait-body scaling relationship (Tang *et al*., 2011; Shingleton and Tang, 2012; Luo *et al*., 2013; Casasa and Moczek, 2018; Okada *et al*., 2019). If there were genetic variation in the growth-sensitivity of individual traits to changes in IIS or TOR signaling, this would generate genetic variation in the slope of the trait-body size morphological scaling relationship. Further, developmental studies suggest that the distribution of slopes (seesaw, speedometer, broomstick) would depend on the locus of genetic variation. For example, changes in the expression of the *Forkhead Transcription Factor* (*FOXO)*, which suppresses growth when nutrition is low but is not active when nutrition is high, generate a broomstick distribution of scaling relationships (Shingleton and Tang, 2012). In contrast, changes in the expression of the *Insulin Receptor* (*InR*), which promotes growth when nutrition is high but is not active when nutrition is low, generate a speedometer distribution (Shingleton and Tang, 2012).

We found that in both males and females, variation in the slope of both the wing-body and leg-body individual scaling relationships correlated most strongly with genetic variation in body size plasticity, rather than size plasticity of the individual traits. This suggests that it is variation in the sensitivity of the body to changes nutrition, independent of the sensitivity of individual traits, that generates variation in the slope of individual scaling relationships. How this is achieved seems paradoxical, since the size of the body ostensibly reflects the collective size of its constituent parts. In *Drosophila*, as with all fully metamorphic insects, the external appendages, such as wings, legs, genitalia, and mouthparts, develop as imaginal discs within the larval body. Pupal size, which we used a proxy for overall body size, is determined by the size of the larva when it stops feeding approximately 24 hours before pupariation. The imaginal discs, however, continue to grow until approximately 24 hours after pupariation (Bryant and Schmidt, 1990). Consequently, the developmental mechanisms that regulate body size are potentially distinct from those that regulate the size of individual traits (Tang *et al*., 2011). Thus, genetic variation in the plasticity of body size and the plasticity of trait size can be distinct from each other. This explanation is supported by the observation that variation in the size plasticities of both the wing and the leg are more tightly correlated with each other than with the body as a whole (Supplementary Table 6). Even though body size plasticity is at least partially independent from trait size plasticities, genetic variation in the former will inevitably result in coordinated changes in the slope of morphological scaling across the traits.

While we have a extensive knowledge of the developmental mechanisms that regulate nutritionally-induced size plasticity of both the body and of individual traits (Nijhout *et al*., 2014), it remains a hypothesis that genetic variation in this plasticity, and by extension in the slope of individual nutritional scaling relationships, lies within these mechanisms. This hypothesis appears to be supported by GWAS studies on thermotolerance and thermally induced body size plasticity, which have identified and functionally validated genes that are involved in the response to environmental change in general and thermal change in particular (Gerken *et al*., 2015; Lafuente *et al*., 2018; Lecheta *et al*., 2020). Consequently, we may expect that genetic variation for nutritionally-induced size variation lies within the developmental pathways involved in the response to nutritional change. The next step is therefore to identify the genes that underlie the observed slope variation in nutritional scaling relationships, and to functionally test their role in regulating the response of body and trait size to variation in developmental nutrition.

## Conclusion

Our data reveal the distribution of previously cryptic individual scaling relationships for wing and leg size against body size in *Drosophila*, and explore their relationship with variation in nutritionally induced plasticity of trait and body size. These data not only provide insight into the genetic architecture of the wing-body and leg-body population scaling relationships, but they also allow us to predict how the population scaling relationship will respond to selection for changes in slope and intercept. Further, future analysis promises to identify the developmental mechanisms that are responsible for the observed genetic variation in individual scaling relationships – the mechanisms that may be targeted by selection to alter population scaling.

## Acknowledgements

This work was made possible through the assistance of undergraduate members of the Shingleton and Frankino labs, who reared and measured the flies used in the study. Additional financial support was provided by the University of Illinois at Chicago.

## Author Contributions

AWS and WAF designed the study; ASW and IMV oversaw the collection of the data; ASW, IMV, WAF and AWS contributed to the data analysis and in preparing the manuscript for publication.

## Data Archiving

All the data and the R scripts used to analyze them are deposited on Dryad.

