## Supplementary for "The Genetic Architecture of Morphological Scaling"

### Supplementary Information

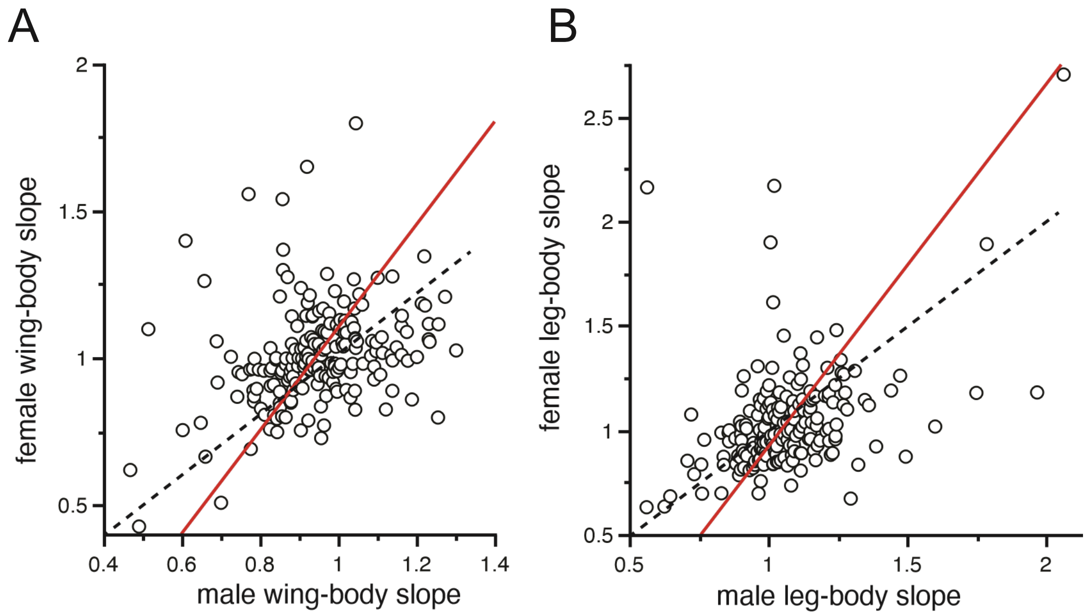


**Supplementary Figure 1:** The relationship between the slope of individual scaling relationships in females and males. (**A**) The relationship between the slope of the wing-body scaling relationship in males versus females is steeper than 1 (solid line, MA slope = 1.76, 95% CI = 1.13-3.07). (**B**) The relationship between the slope of the leg-body scaling relationship in males versus females is also steeper than 1 (solid line, MA slope = 1.74, 95% CI = 1.28-2.49). Dotted line is where female slope = male slope.

**Supplementary Table 1:** Effect of 'Block' on trait size.

| Trait[a] | Block SS | Residual SS | Block DF | Residual DF | *F* | *P*[a] |
|---|---|---|---|---|---|---|
| Wing | 12.97 | 867.8 | 8 | 22349 | 41.744 | <0.001 |
| Leg | 28.01 | 772.76 | 8 | 22349 | 101.26 | <0.001 |
| Pupa | 41.89 | 788.29 | 8 | 22349 | 148.46 | <0.001 |

[a] Data were fit with the model $T = B$, where $T$ is trait size and $B$ is block.

**Supplementary Table 2:** Effect of sex on the wing-body individual morphological scaling relationship.

| Effect | Effect Size[a] | SE | DF | t | *P* |
|---|---|---|---|---|---|
| (Intercept) | 0.0573 | 0.00427 | 198 | 13.411 | <0.001*** |
| Sex (male) | -0.1060 | 0.00103 | 22100 | -102.752 | <0.001*** |
| Body Size | 0.8884 | 0.00819 | 240 | 107.945 | <0.001*** |
| Sex(male)*Body Size | -0.0550 | 0.00553 | 22170 | -9.947 | <0.001*** |

[a] The effect size is from the mixed linear model $T = S + B + S*B + L$, where $T$ is wing size, $S$ is sex, $B$ is body size, $S*B$ is the sex-by-body size interaction (where the wing-body relationship varies by sex) and $L$ is lineage (random factor). Parameter values are zero unless specified.

[ns] *P* value > 0.05, **P* value < 0.05, **P* value < 0.01, *** *P* value < 0.001.

**Supplementary Table 3:** Effect of sex on the leg-body individual morphological scaling relationship.

| Effect | Effect Size[a] | SE | DF | t | *P* |
|---|---|---|---|---|---|
| (Intercept) | 0.0004 | 0.0043 | 200 | -0.0930 | 0.9261 [ns] |
| Sex (male) | 0.0156 | 0.0012 | 22100 | 9.3450 | <0.001*** |
| Body Size | 0.8842 | 0.0098 | 230 | 86.5200 | <0.001*** |
| Sex(male)*Body Size | 0.0148 | 0.0055 | 22170 | 2.2300 | 0.0258* |

[a] The effect size is from the mixed linear model $T = S + B + S*B + L$, where $T$ is leg size, $S$ is sex, $B$ is body size, $S*B$ is the sex-by-body size interaction (where the wing-body relationship varies by sex) and $L$ is lineage (random factor). Parameter values are zero unless specified.

[ns] *P* value > 0.05, **P* value < 0.05, **P* value < 0.01, *** *P* value < 0.001.

**Supplementary Table 4:** Likelihood ratio test that MA slopes vary among lineages

| Trait | Sex | Likelihood Ratio[a] | DF | *P* |
|---|---|---|---|---|
| Wing | Male | 603.4 | 193 | <0.001*** |
| | Female | 644.7 | 194 | <0.001*** |
| Leg | Male | 520.3 | 193 | <0.001*** |
| | Female | 544.8 | 194 | <0.001*** |

[a] Likelihood ratio test of the null hull hypothesis that the slopes are equal.

*** P value < 0.001.

**Supplementary Table 5:** Effect of including random variation in slope on the fit of the relationship between wing or leg and body size in males and females

| Trait | Sex | Model[a] | DF[b] | AIC[c] | BIC[c] | Likelihood Ratio[c] | *P*[d] | Variance[e] |
|---|---|---|---|---|---|---|---|---|
| Wing | Male | Model 1: $W_{jk}=B_j+B_j*L_k$ | 7 | -27647.7 | -27603.7 | 471.9934 | <0.001*** | 0.012 (0.009-0.016) |
| | | Model 2: $W_{jk}=B_j+L_k$ | 5 | -27179.7 | -27150.4 | | | |
| | Females | Model 1: $W_{jk}=B_j+B_j*L_k$ | 7 | -24694.6 | -24650.8 | 468.504 | <0.001*** | 0..012 (0.008-0.015) |
| | | Model 2: $W_{jk}=B_j+L_k$ | 5 | -24230.1 | -24200.9 | | | |
| Leg | Male | Model 1: $W_{jk}=B_j+B_j*L_k$ | 7 | -22331.9 | -22287.8 | 353.4541 | <0.001*** | 0.015 (0.011-0.020) |
| | | Model 2: $W_{jk}=B_j+L_k$ | 5 | -21982.4 | -21953.1 | | | |
| | Females | Model 1: $W_{jk}=B_j+B_j*L_k$ | 7 | -21098.6 | -21054.8 | 296.408 | <0.001*** | 0.013 (0.009-0.017) |
| | | Model 2: $W_{jk}=B_j+L_k$ | 5 | -20806.2 | -20777 | | | |

[a] *T* is trait size, *B* is body size, and L is lineage (random effect). The models differ by having random slopes and intercepts (model 1) versus random intercepts (model 2) among lineages.

[b] Estimated degrees of freedom for each model.

[c] AIC, BIC, and likelihood ratio calculated using ML fit.

[d] *P*-value of calculated by parametric bootstrapping using ML fit with 1000 replicates. [ns] *P* value > 0.05, **P* value < 0.05, **P* value < 0.01, *** *P* value < 0.001.

[e] Variance of slope among lineages calculated using Bayesian fit, with 95% confidence interval.

**Supplementary Table 6:** Variances (diagonal), Covariances (below diagonal) and Pearson's Correlations (above diagonal) of trait size plasticity, independent of body size plasticity, within and between sexes [a]

| | | male | | female | |
|---|---|---|---|---|---|
| | | wing | leg | wing | leg |
| male | wing | **0.0050** | 0.66 | 0.35 | 0.20 |
| | leg | 0.0039 | **0.0072** | 0.27 | 0.33 |
| female | wing | 0.0019 | 0.0017 | **0.0057** | 0.59 |
| | leg | 0.0012 | 0.0024 | 0.0039 | **0.0076** |

[a] The darker color the higher the covariance/correlation. All correlations are significant at *P*<0.0001.

**Supplementary Table 7:** $R^2$ of linear relationships between wing-, leg- and body-plasticity and the slope of the wing/leg-body individual scaling relationship, among lineages.

| **Sex** | **Trait-plasticity (*x*-axis)** | **Scaling Relationship (*y*-axis)** | **$R^2$** | **95% Confidence Interval** |
|---|---|---|---|---|
| Male | Wing | Wing-Body | 0.087*** | 0.013–0.162 |
| | Body | Wing-Body | 0.123*** | 0.038–0.208 |
| Female | Wing | Wing-Body | 0.0137$^{ns}$ | -0.018–0.046 |
| | Body | Wing-Body | 0.286*** | 0.180–0.391 |
| Male | Leg | Leg-Body | 0.100*** | 0.021–0.178 |
| | Body | Leg-Body | 0.106*** | 0.025–0.186 |
| Female | Leg | Leg-Body | 0.009$^{ns}$ | -0.017–0.035 |
| | Body | Leg-Body | 0.251*** | 0.147–0.355 |

$^{ns}$ *P* value > 0.05, **P* value < 0.05, ***P* value < 0.01, *** *P* value < 0.001.
